# Multiple sclerosis-associated HLA demarcates EBV-specific CD8^+^ T cells with an exhausted and brain residency phenotype

**DOI:** 10.1101/2025.06.27.661912

**Authors:** Sanne Reijm, Ana M. Marques, Jasper Rip, Cato E.A. Corsten, Annet F. Wierenga-Wolf, Harm de Wit, Marie-José Melief, Yifan van Hasselt, Jamie van Langelaar, Rinze Neuteboom, Beatrijs H.A. Wokke, Yvonne M. Mueller, Joost Smolders, Marvin M. van Luijn

## Abstract

Specific HLA class I subtypes as well as circulating antibodies against Epstein-Barr virus (EBV) have been associated with increased susceptibility to develop multiple sclerosis (MS). It is currently unclear whether and how these risk factors are functionally related. Here, we assessed the impact of MS risk allele HLA-B7 and protective allele HLA-A2 on the effector phenotype of EBV-specific CD8^+^ T cells and if this corresponds with anti-EBV IgG titers. For this, we selected HLA-A2^+^ and/or HLA-B7^+^ donors with and without MS and analyzed HLA-restricted, EBV-specific CD8^+^ T cells with different fine-specificities using 38 color-based spectral flow cytometry. In contrast to HLA-A2-restricted CD8^+^ T cells, HLA-B7-restricted CD8^+^ T cells showed a confined response to EBV, which was mainly directed against latent peptide EBNA3A(379-387). These HLA-B7-restricted EBV-specific CD8^+^ T cells expressed higher levels of CNS residency markers CXCR3 and CD20 and were most abundant in fresh single-cell suspensions derived from different post-mortem CNS compartments of an HLA-A2^+^B7^+^ MS donor. Moreover, HLA-B7- restricted EBV-specific CD8^+^ T cells displayed a more exhausted phenotype (PD- 1^+^CD244^+^CD160^+^KLRG1^+^TIGIT^+^) that stood out in the blood from people with MS (pwMS). Accordingly, in the blood, increased IgG titers against EBV were found for pwMS carrying HLA- B7 but lacking HLA-A2, which seemed to coincide with HLA-B7-restricted EBV-specific CD8^+^ T cells having such an effector phenotype. These data support a model in which the confined response against EBV makes circulating HLA-B7-restricted CD8^+^ T cells less able to control EBV and more prone to infiltrate the CNS in pwMS.

## Introduction

Multiple sclerosis (MS) is a chronic inflammatory disease leading to neurodegeneration in the central nervous system (CNS). The cause of MS is unknown, but genetic predisposition together with environmental influences are considered to be the disease trigger. Large genome-wide association studies have identified 233 genetic risk variants, of which 32 are located in the HLA region (*1*). This region is the by far strongest associated with MS. Especially the HLA class I locus contains alleles that can either have an independent risk or protective effect, of which the most common are HLA-B7 and HLA-A2, respectively (*2–5*). The major environmental susceptibility factor for MS is an infection with the Epstein-Barr virus (EBV), as now proven by a required specific antibody response prior to onset. An infection with the cytomegalovirus (CMV), another persistent herpesvirus, does not increase but might even decrease this risk (*6*). Furthermore, it has been shown that antibody levels against EBV are increased in people with MS (pwMS) compared to the healthy population (*7*). Together, this suggests that in conjunction with an increased seroprevalence, the cellular immune response against EBV is differentially regulated because of the presence of MS risk versus protective HLA class I alleles.

Immune infiltration in the brain is a hallmark of MS attacks and the main difference between the CNS of people with and without MS is the presence of B cells (*8*). Since CD8^+^ T cells are important players in controlling viral infections and can recognize EBV-derived peptides in MS-associated risk or protective HLA class I alleles presented by infected B cells, these cells might be involved in the aberrant immune response against EBV in MS. CD8^+^ T cells are the major fraction of T cells populating the brain of pwMS (*9*), aberrant recruitment of CD8^+^ T cells has been described in MS cerebrospinal fluid (CSF) (*10*), and CD8^+^ T cells directed against EBV antigens have been identified in the CSF of pwMS (*11*). Lower expression of pro-inflammatory and cytolytic proteins by EBV-specific CD8^+^ T cells after antigen stimulation has been described for the HLA-B7 risk allele in pwMS compared to healthy people (*12*). Moreover, EBV viral load was found to be higher in HLA-B7^+^ compared to HLA-A2^+^ pwMS (*13*). Together, this suggests that HLA-B7 and/or HLA-A2 carriage determines the ability of CD8^+^ T cells to control the immune response against EBV inside and/or outside the CNS in people who develop MS.

Little is known about how fine-specificities to EBV distinctively shape the effector phenotype of CD8^+^ T cells and their potential to infiltrate the CNS in the context of protective and risk HLA alleles. Together with a potential role in EBV seroprevalence, deeper analysis of such T-cell subsets could improve the understanding of MS susceptibility and support the development of EBV-specific T-cell therapies. Therefore, we performed an in-depth phenotypical characterization of HLA-B7- and HLA-A2-restricted CD8^+^ T cells specific for different EBV and CMV epitopes from the blood of healthy individuals and treatment-naive as well as natalizumab-treated pwMS. The treatment with natalizumab (anti-VLA-4) allowed us to analyze CNS-homing CD8^+^ T cells that are trapped in the blood. Furthermore, we assessed the recruitment and effector phenotype of fine-specific CD8^+^ T cells in different post-mortem CNS compartments from an MS donor *ex vivo*.

## Results

### The number of EBV epitopes is confined for HLA-B7-restricted CD8^+^ T cells

Since the susceptibility to develop MS is increased by an infection with EBV and the presence of HLA risk allele B7 and decreased by the presence of HLA protective allele A2, we first investigated the effect of HLA-type on the abundance/frequencies of EBV-specific CD8^+^ T cells in healthy donors (HD). CMV-specific CD8^+^ T cells served as antigen-specific controls. Antigen-specific cells were detected by a combinatorial staining method with HLA class I tetramers (Fig. S1A), resulting in simultaneous detection of CD8^+^ T cells recognizing 10 different HLA-A2-restricted or 8 different HLA-B7-restricted EBV- and CMV immunodominant epitopes. A representative staining of HLA-A2- and HLA-B7 restricted CD8^+^ T cells from HD blood is shown in Fig. S1B and Fig. S1C, respectively. We determined the frequencies of EBV- and CMV-specific CD8^+^ T cells specific for immunodominant epitopes presented in HLA-A2 or HLA-B7. For EBV, peptides derived from both lytic and latent proteins were included. For HLA-A2, various lytic and latent epitopes were recognized by the EBV-specific CD8^+^ T cells (Fig. 1A), while the recognition of EBV-derived peptides in HLA-B7 was restricted to mainly one latent epitope, EBNA3A(379-387) (Fig. 1B). For subsequent analysis, CD8^+^ T cells were distinguished based their specificity to EBV latent or lytic peptides. More EBV latent than lytic peptides were recognized by HLA-B7-restricted CD8^+^ T cells, which was not the case for EBV-specific CD8^+^ T cells restricted to HLA-A2 (Fig. 1C). We found no significant differences in total EBV- or CMV-specific CD8^+^ T-cell frequencies between HLA-A2 and HLA-B7 carriers (Fig.1D).

**Figure 1.**
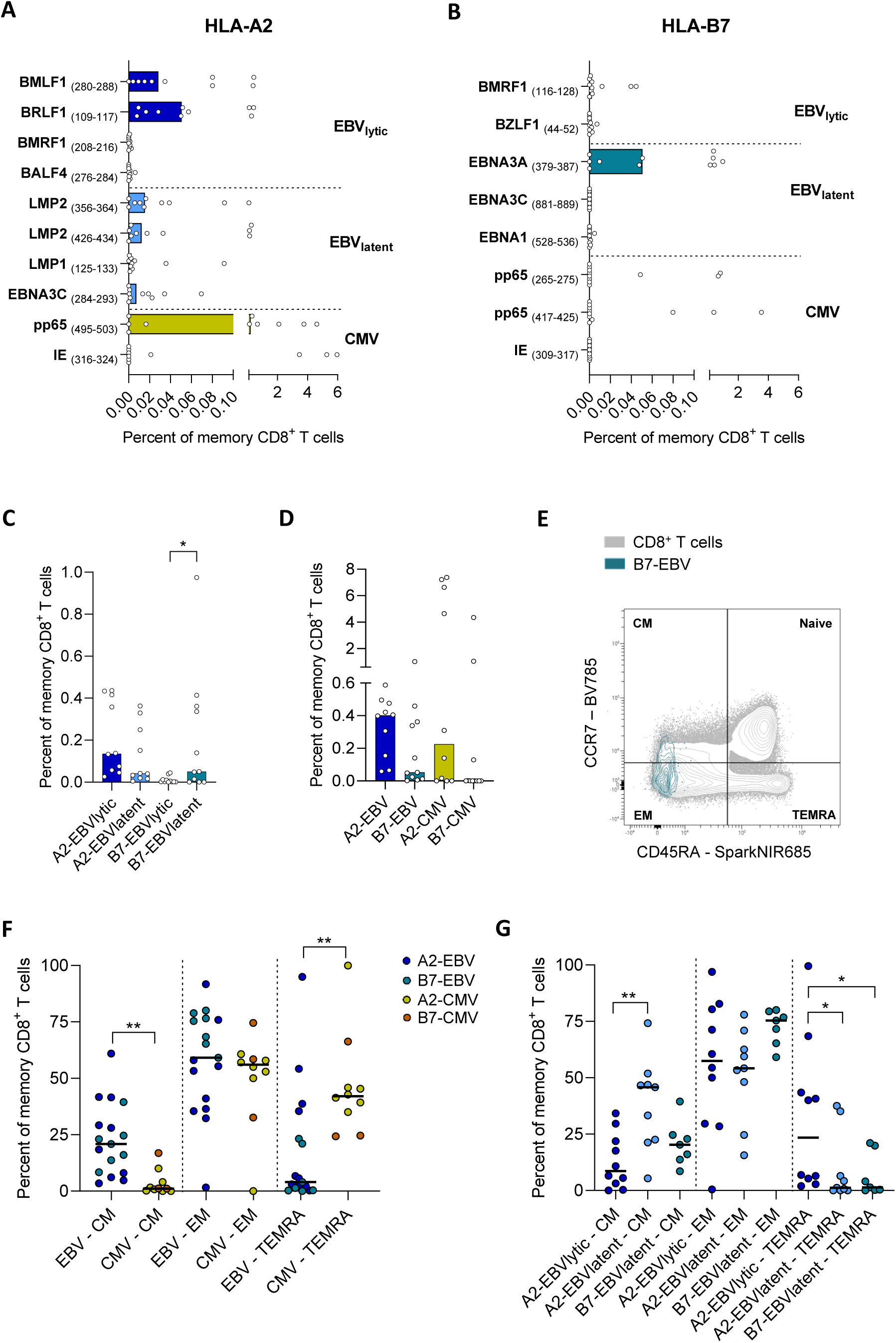
Frequencies and differentiation of EBV- and CMV-specific CD8+ T cells in HD. Percentage of EBV- and CMV-specific CD8+ T cells of total memory CD8+ T cells per immunodominant peptide restricted to HLA-A2 (A) or HLA-B7 (B). (C) Percentages of EBV-specific CD8+ T cells of total memory CD8+ T cells restricted to latent or lytic peptides. Wilcoxon test was used for statistical testing. (D) Percentage of total EBV- and CMV-specific CD8+ T cells of total memory CD8+ T cells. Mann-Whitney test was used for statistical testing. (E) representative example of the gating strategy of the differentiation states. (F) Percentage of total EBV- and CMV-specific CD8+ T cells of total memory CD8+ T cells per differentiation state. Mann-Whitney test was used for statistical testing. (G) Percentages of EBV-specific CD8+ T cells of total memory CD8+ T cells restricted to latent or lytic peptides per differentiation state. Kruskal-Wallis test with Dunn posthoc was used for statistical testing. Every dot represents one individual. Bars or lines represent medians. *p<0.05 ** p<0.005 ***p<0.0005 **** p<0.0001.

### Antigen-specificity and HLA distinctively shapes the EBV-specific CD8^+^ T-cell memory state

Next, we investigated how the presence of HLA-B7 and HLA-A2 is associated with the memory state of EBV- and CMV-specific CD8^+^ T cells based on the expression of CCR7 and CD45RA (Fig. 1E). CMV-specific cells especially showed an effector memory (EM) or terminally differentiated effector memory (TEMRA) phenotype, while EBV-specific T cells were also found within the central memory (CM) compartment (Fig. 1F). Within the pool of HLA-A2-restricted EBV-specific CD8^+^ T cells, those specific for EBV lytic peptides were mostly EM and TEMRA and those specific for EBV latent peptides were mostly CM (Fig. 1G). This could be explained by the nature of the immunodominant peptide, as lytic proteins are expressed during the replication phase and therefore antigen exposure will be different from the proteins expressed during the latent phase of an EBV infection. However, HLA-B7-restricted CD8^+^ T cells specific for EBV latent peptide displayed a memory phenotype that was similar to HLA-A2-restricted CD8^+^ T cells specific for EBV lytic peptides and tended to be more EM (Fig. 1G).

### Natalizumab restores the decrease of EBV-specific CD8^+^ T_EM_ cells in MS

To study whether HLA-type differentially impact the virus-specific T-cells in the context of MS, we investigated frequencies and differentiation state of EBV- and CMV-specific CD8^+^ T cells from pwMS. No differences in total frequencies of EBV- and CMV-specific memory CD8^+^ T cells were found between the untreated MS (UT-MS) and natalizumab-treated MS (NTZ-MS) group. Irrespective of natalizumab treatment, the frequencies of EBV-specific CD8^+^ T cells tended to be lower for those restricted to HLA-A2 and higher for those restricted to HLA-B7 in pwMS compared to HD (represented by the dotted line), although these differences were not significant (Fig. 2A). The frequencies of HLA-B7-restricted CMV-specific CD8^+^ T cells were only slightly higher in pwMS (Fig. 2B). Within the pool of EBV-specific CD8^+^ T cells, these differences were driven by lytic peptide for those restricted to HLA-A2 and latent peptide for those restricted to HLA-B7 (Fig. 2C). The frequencies of all separate fine-specific CD8^+^ T cells are shown in Fig. S2. When comparing the percentages of EBV- and CMV-specific CD8^+^ T cells in HLA-A2^+^B7^+^ positive donors in a paired manner, HLA-B7-restricted cells were more abundant than HLA-A2- restricted cells (Fig. 2D). This was significant for the CMV-specific cells and a similar trend was seen for EBV latent peptide-specific cells (Fig. 2D). Next, we compared the memory state of EBV- and CMV-specific CD8^+^ T cells between HD and pwMS. For UT-MS, a decrease was detected for EBV- and not for CMV-specific CD8^+^ T_EM_ cells when compared to HD, which was restored in the NTZ-MS group (Fig. 2E). This was especially seen for EM CD8^+^ T_EM_ cells specific for EBV latent peptide and irrespective of restriction to HLA-B7 or HLA-A2. (Fig. 2F). This suggests that EBV-specific CD8^+^ T_EM_ cells have an increased potential to traffic towards the CNS.

**Figure 2.**
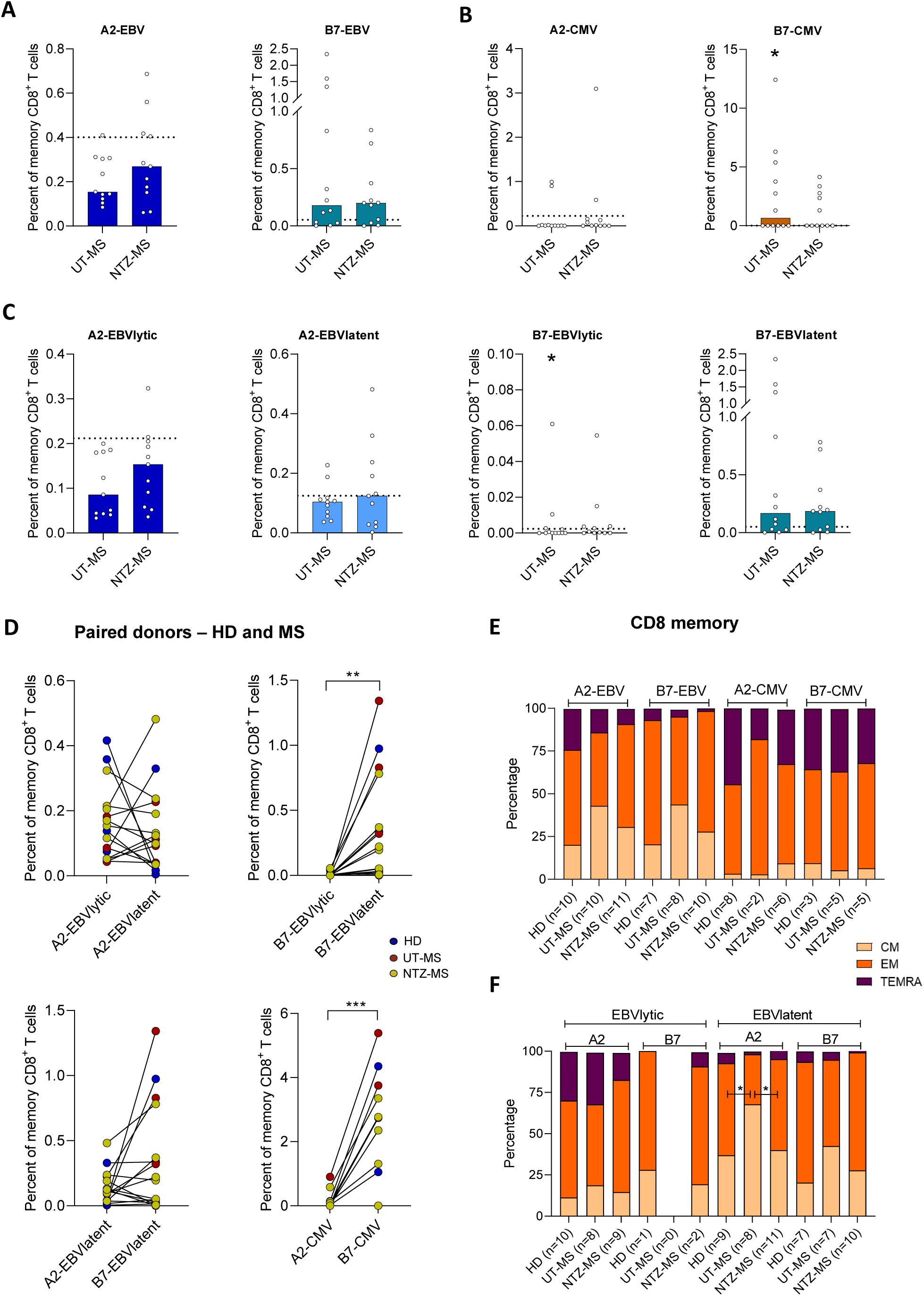
Frequencies and differentiation of EBV- and CMV-specific CD8+ T cells in MS. Percentage of antigen-specific CD8+ T cells of total memory CD8+ T cells for MS cohorts for total EBV (A), total CMV (B) or EBV stratified for latent and lytic peptides (C). Mann-Whitney test was used for statistical testing. (D) Percentage of EBV- or CMV-specific CD8+ T cells of total memory CD8+ T cells in a paired manner. Wilcoxon test was used for statistical testing. (E) Differentiation stages of EBV- and CMV-specific CD8+ T cells for 3 cohorts. Kruskal-Wallis test with Dunn posthoc was used for statistical testing. (F) Differentiation stages of EBV-specific CD8+ T cells restricted to latent or lytic peptides for 3 cohorts. Kruskal-Wallis test with Dunn posthoc was used for statistical testing. Every dot represents one individual. Bars represent medians. Dotted line in A, B and C represents median of HD. *p<0.05 ** p<0.005 ***p<0.0005 **** p<0.0001.

### CD20^dim^ expression characterizes HLA-B7-restricted EBV-specific CD8^+^ T cells

To further elaborate on the CNS-homing potential of EBV-specific memory CD8^+^ T cells, we developed a spectral flow cytometry panel and evaluated tissue-homing and -residency associated markers (Fig. 3A-B). No differences in expression level of these markers were found between the UT-MS and NTZ-MS group. cohorts, so these data were combined for further analysis as a whole group. Furthermore, HLA-B7-restricted EBV lytic peptide-specific CD8^+^ T cells were excluded from the analysis since very few cells could be detected as shown in Fig. 1 and 2. The expression level of CXCR3 was increased on EBV-specific compared to non-EBV/CMV CD8^+^ memory T cells and CMV-specific CD8^+^ memory T cells from HD and pwMS (Fig. 3C), which can be linked to the high expression of this marker on T cells residing in the CNS (*9*) and high concentrations of its ligand CXCL10 in CSF (*14*). This high CXCR3 expression on EBV-specific CD8^+^ T cells was not affected by restriction to HLA-B7 and HLA-A2 or specific for MS (Fig. 3D). Furthermore, CD20 is enriched on circulating CD8^+^ T cells has been described in MS and is even further upregulated on T cells residing in CNS tissues (*15, 16*). Interestingly, CD20 expression was increased on EBV-specific compared to non-EBV/CMV CD8^+^ memory T cells and CMV-specific CD8^+^ T cells, which was found for both the HD and MS group (Fig. 3E). When looking into fine specificity and HLA restriction, CD20 expression was most pronounced on HLA-B7- compared to HLA-A2-restricted EBV-specific CD8^+^ T cells in the MS group (Fig. 3F). For all other tissue-homing or residency associated markers, no significant differences were found between subgroups. Together, these data suggest that particularly HLA-B7-restricted EBV-specific CD8^+^ T cells are better equipped to enter and reside in the CNS.

**Figure 3.**
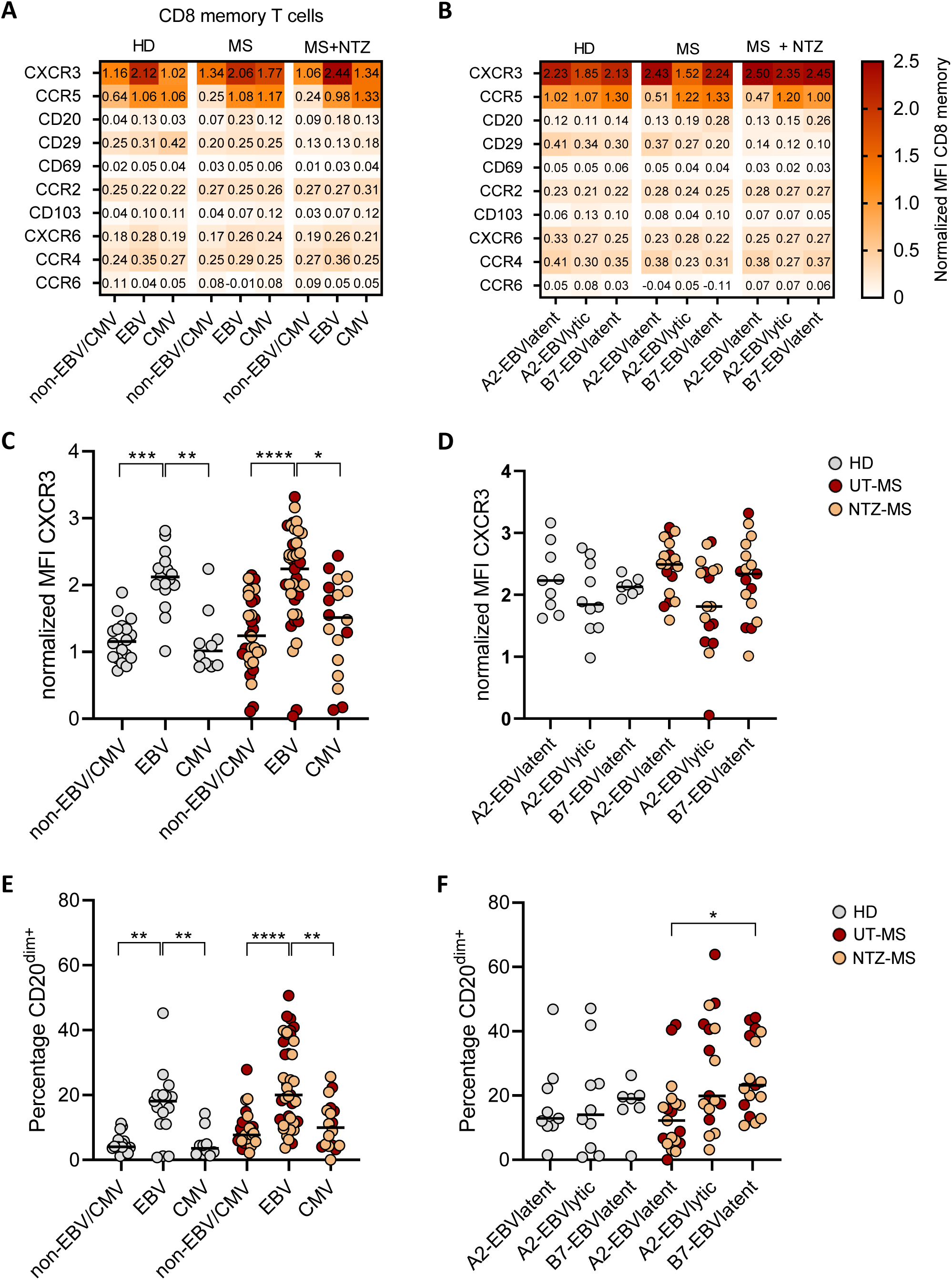
Brain homing and brain-residency associated markers on EBV- and CMV-specific CD8+ T cells in HD and MS. Heatmap of median protein expression in normalized median fluorescence intensity (MFI) for total CD8+ memory T cells, EBV- and CMV-specific CD8+ memory T cells (A) or for EBV-specific CD8+ memory T cells restricted to latent or lytic peptides (B). CXCR3 expression for total CD8+ memory T cells, EBV- and CMV-specific CD8+ memory T cells (C) or for EBV-specific CD8+ memory T cells restricted to latent or lytic peptides (D). CD20dim expression for total CD8+ memory T cells, EBV- and CMV-specific CD8+ memory T cells (E) or for EBV-specific CD8+ memory T cells restricted to latent or lytic peptides (F). Kruskal-Wallis test with Dunn posthoc was used for statistical testing. Every dot represents one individual. Black lines represent medians. *p<0.05 ** p<0.005 ***p<0.0005 **** p<0.0001.

### HLA-B7-restricted EBV-specific CD8^+^ T cells have an exhausted phenotype in MS

To further characterize the effector phenotype of EBV-specific CD8^+^ T cells, we compared surface levels of activation, co-stimulatory, co-inhibitory and cytotoxicity-related markers between the HLA and antigen-dependent subgroups. No differences in expression were found between the UT-MS and NTZ-MS groups, so these data were combined in further analysis. EBV-specific CD8^+^ T cells were distinguished from CMV-specific CD8^+^ T cells by a higher expression of CD28, TIGIT and CD127 (IL-7R) and a lower expression of GPR56, CX3CR1 and KLRG1 (Fig. 4A and Fig. S3), which is in line with previous studies (*17*). Co-inhibitory receptors CD244, PD-1 and CD160 were all higher expressed on both EBV- and CMV-specific compared to non-EBV/CMV CD8^+^ memory T cells. These differences were found for both the HD and MS group (Fig. S3).The differences in expression of co-stimulatory and co-inhibitory markers stood out when comparing HLA-A2-restricted CD8^+^ T cells specific for EBV latent and lytic peptide with HLA-B7-restricted CD8^+^ T cells specific for EBV latent peptide (Fig. 4B). Lower expression of co-stimulatory marker CD28 and higher expression of co-inhibitory receptors CD244, CD160, KLRG1 and TIGIT was found for HLA-B7- versus HLA-A2-restricted CD8^+^ T cells specific for EBV latent peptide (Fig. S4). These features were similar to HLA-A2-restricted CD8^+^ T cells specific for EBV lytic peptides, except that TIGIT was higher expressed on HLA-B7-restricted EBV latent peptide-specific CD8^+^ T cells (Fig. S4F). As co-expression of co-inhibitory receptors is one of the hallmarks of T-cell exhaustion, we subsequently evaluated the co-expression of PD-1, CD244, CD160, KLRG1 and TIGIT. A representative example for the gating of each co-inhibitory receptor is shown in Fig. 4C. In comparison to both non-EBV/CMV- and CMV-specific CD8^+^ T cells, this co-expression profile was especially found for EBV-specific CD8^+^ T cells, in particular for those restricted to HLA-A2-bound lytic peptides or HLA-B7-bound latent peptides. Co-expression of these co-inhibitory receptors was most distinctive in pwMS and tended to be highest in HLA-B7- restricted CD8^+^ T cells (Fig. 4D). These differences in co-expression on HLA-restricted and fine-specific CD8^+^ T cells were also observed in a paired manner within the same individuals (Fig. 4E). Among the non-EBV/CMV-specific CD8^+^ memory T cells, this co-expression was increased on EM and TEMRA compared to CM cells (Fig. 4F), which could influence our results since the A2-EBVlatent specific cells included more CM cells. However, when we stratified for memory state, the co-expression profile was still more pronounced for HLA-B7- than HLA-A2-restricted CD8^+^ T cells specific for EBV latent peptide and was mostly seen within the EM compartment (Fig. 4G). Together, these results suggest that HLA-B7-restricted EBV-specific T cells are more exhausted and might be less able to control EBV even during the latent phase of the infection.

**Figure 4.**
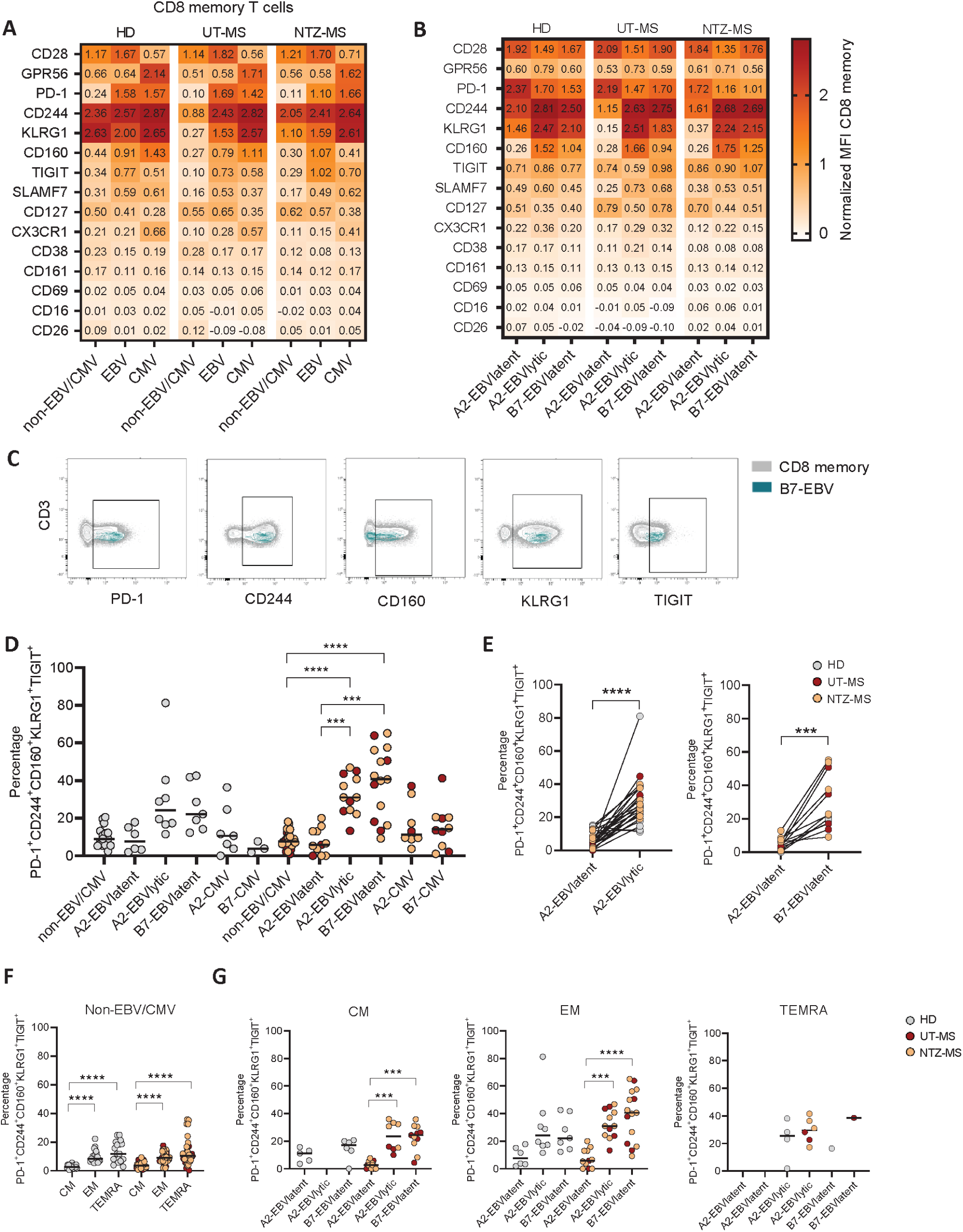
Activation-associated markers on EBV- and CMV-specific CD8+ T cells in HD and MS. Heatmap of median protein expression in normalized median fluorescence intensity (MFI) for total CD8+ memory T cells, EBV- and CMV-specific CD8+ memory T cells (A) or for EBV-specific CD8+ memory T cells restricted to latent or lytic peptides (B). (C) Representative example of gating of co-inhibitory receptors. (D) Exhaustion phenotype on CMV-specific CD8+ memory T cells and EBV-specific CD8+ memory T cells restricted to latent or lytic peptides. Kruskal-Wallis test with Dunn posthoc was used for statistical testing. (E) Exhaustion phenotype on EBV-specific CD8+ memory T cells restricted to latent or lytic peptides in a paired manner. Wilcoxon test was used for statistical testing. (F) Exhaustion phenotype on non-EBV/CMV CD8+ memory T cells. Kruskal Wallis test with Dunn posthoc was used for statistical testing. (G) Exhaustion phenotype on EBV-specific CD8+ memory T cells restricted to latent or lytic peptides stratified by differentiation state. Kruskal Wallis test with Dunn posthoc was used for statistical testing. Every dot represents one individual. Black lines represent medians. *p<0.05 ** p<0.005 ***p<0.0005 **** p<0.0001.

### MS brain-compartments contain HLA-B7-restricted EBV-specific CD8^+^ T cells

To explore how HLA restriction determines the presence of EBV-specific CD8^+^ T cells in the MS CNS, we isolated lymphocytes from post-mortem cerebrospinal fluid (CSF), leptomeninges (LM), normal appearing white matter (NAWM) and affected white matter (lesion) of an MS donor carrying both HLA-B7 and HLA-A2. Both EBV- and CMV-specific CD8^+^ T cells could be detected in the blood and the majority of CNS compartments, but their distribution was dependent on the type of HLA restriction (Fig. 5A-B, Fig. S5). Notably, within the CNS compartments, HLA- B7-restricted EBV-specific CD8^+^ T cells were most abundantly found, which was vice versa for CMV-specific CD8*^+^* T cells (Fig. 5C). In line with our findings in pre-mortem blood and in contrast to CMV-specific counterparts, EBV-specific CD8^+^ T cells had predominantly an EM phenotype. We did not find any differences in memory state of EBV- or CMV-specific CD8^+^ T cells between the blood and different CNS compartments (Fig. 5D). As expected, tissue-homing and residency associated markers were upregulated on total CD8^+^ memory T cells in all CNS compartments as compared to blood (Fig. S6A). Interestingly, HLA-B7-restricted EBV-specific CD8^+^ T cells expressed the highest level of these markers, especially in the CSF (Fig. S6A). Within all compartments, tissue residency-associated markers CD20 and CXCR6 were selectively abundant on HLA-B7-restricted EBV-specific CD8^+^ T cells, which was the highest in LM and affected white matter (Fig. 5E, Fig. S6A). In addition, the pronounced co-expression profile of co-inhibitory markers as observed for HLA-B7-restricted EBV-specific CD8^+^ T cells was seen in all compartments, but, importantly, less pronounced within the CNS than the blood (Fig. 5F, Fig. S6B). Of the CNS compartments, this co-expression was the highest in the LM, in which most B cells are found post-mortem in MS (*18*). In conclusion, in this MS case, predominantly HLA-B7- restricted EBV-specific EM CD8^+^ T cells have recruited to the CNS, of which a minority show an exhausted phenotype.

**Figure 5.**
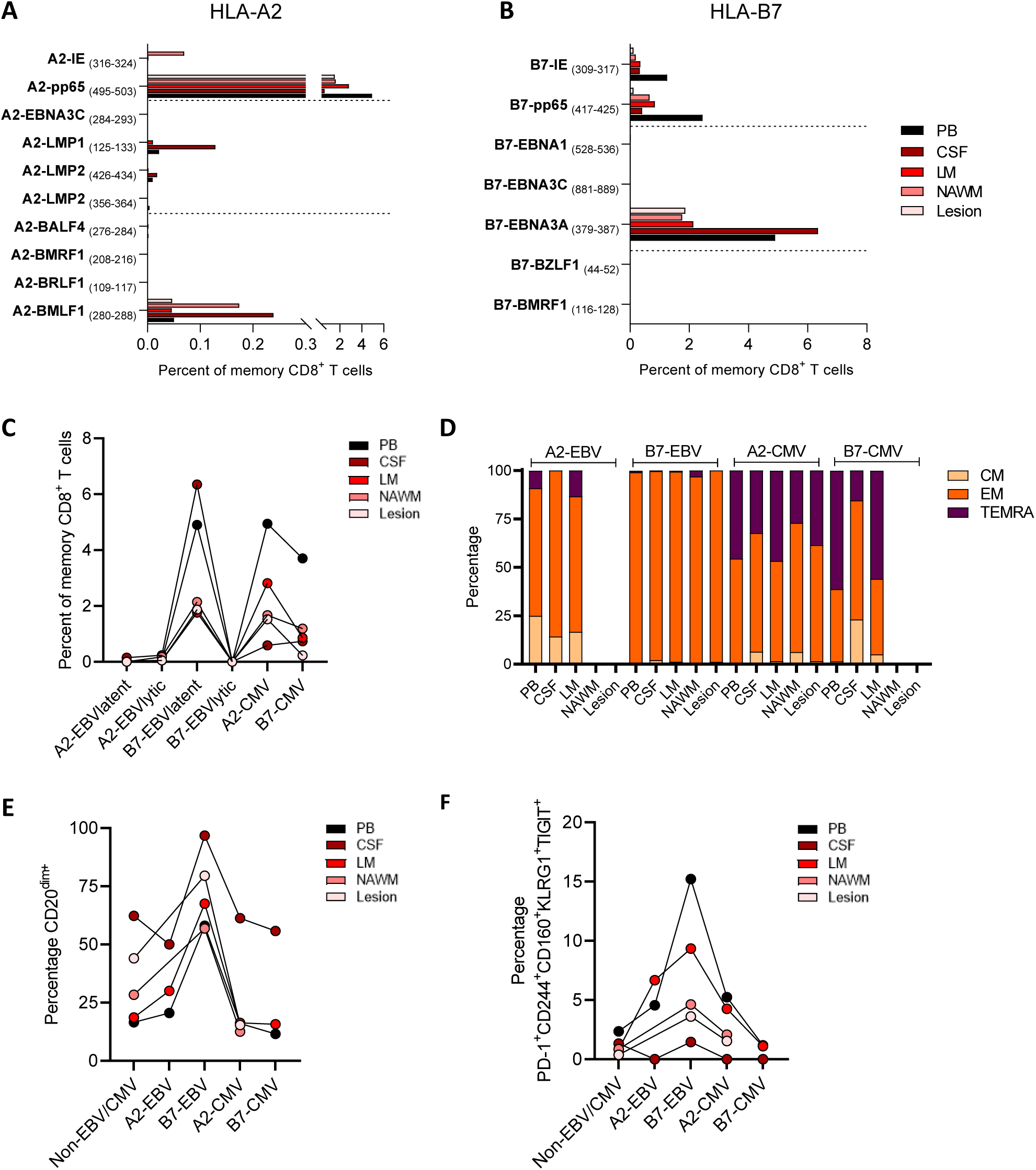
EBV- and CMV-specific CD8+ T cells in CNS compartments. Percentage of EBV- and CMV-specific CD8+ T cells of total memory CD8+ T cells in blood and CNS compartments per immunodominant peptide restricted to HLA-A2 (A) or HLA-B7 (B). (C) Percentages of EBV-specific CD8+ T cells of total memory CD8+ T cells restricted to latent or lytic peptides in blood and CNS compartments. (D) Differentiation states of EBV- and CMV-specific CD8+ T cells in blood and CNS compartments. (E) Expression of CD20dim of EBV- and CMV-specific CD8+ T cells in blood and CNS compartments. (F) Exhaustion phenotype of EBV- and CMV-specific CD8+ T cells in blood and CNS compartments. Peripheral blood (PB), Cerebrospinal fluid (CSF), leptomeninges (LM), Normal appearing white matter (NAWM).

### EBV-specific CD8^+^ T cell exhaustion phenotype correlates with anti-EBNA1 antibodies

To explore how HLA-B7 and/or HLA-A2 carriage is associated with differential control of EBV, we measured serum antibody titers against EBV and CMV in a cohort of early-stage MS (n=78). Antibody titers against EBV nuclear antigen 1 (EBNA1) and viral capsid antigen (VCA) were significantly higher in HLA-B7^+^HLA-A2^-^compared to HLA-B7^-^HLA-A2^+^ pwMS (Fig. 6AB). This was not found for early antigen (EA)- and CMV-specific antibodies (Fig. 6C-D). After stratification for MS risk allele HLA-DRB1*15:01, which is in linkage disequilibrium with HLA- B7 (*19*), we found that only anti-VCA and anti-CMV antibody titers were significantly elevated in pwMS carrying HLA-DRB1*15:01 (Fig. S7). This eliminated HLA-DRB1*15:01 as a confounder for our observed association between HLA-B7 and/or HLA-A2 carriage and anti- EBNA1 antibody levels. To assess whether anti-EBNA1 antibody levels correspond to the effector phenotype of EBV-specific EM CD8^+^ T cells, we were able to measure our earlier observed co-expression profile for 13 pwMS and found a weak correlation (Fig. 6E). However, this correlation became stronger when selecting for HLA-A2-restricted EBV lytic peptide and HLA-B7-restricted EBV latent peptide-specific EM CD8^+^ T cells (Fig. 6F), for which the expression profile was seen the most (Fig. 4D). These data support the hypothesis that CD8^+^ T cells restricted to HLA-B7 are more exhausted and therefore less capable of controlling an EBV infection, therefore increasing the risk of an increased EBV seroprevalence in MS (Fig. 7).

**Figure 6.**
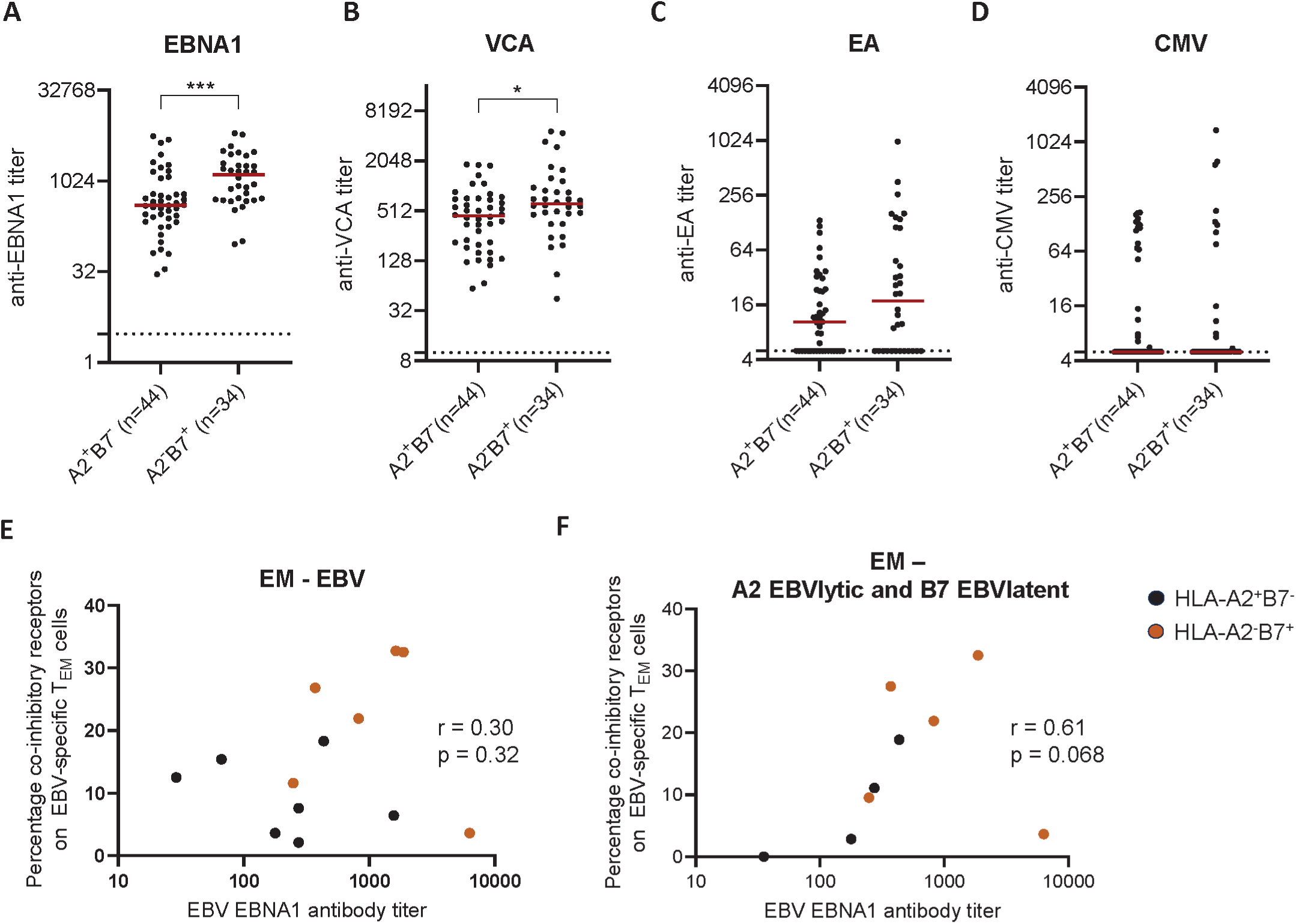
Correlation EBV antibody titers and exhaustion phenotype. Antibody titers of HLA-A2+B7- or HLA-A2-B7+ untreated persons with RRMS from the PROUD cohort. (A) EBNA1 (B) Viral capsid antigen (VCA) (C) Early antigen (EA) (D) CMV. Unpaired T test was used for statistical testing. (E) Correlation of the percentage exhaustion phenotype on EBV-specific TEM cells and EBNA1 antibody titer. UT-MS and NTZ-MS were included. Spearman correlation was used for statistical testing. (F) As in E, but only A2-EBVlytic and B7-EBV latent specific cells are shown.

**Figure 7.**
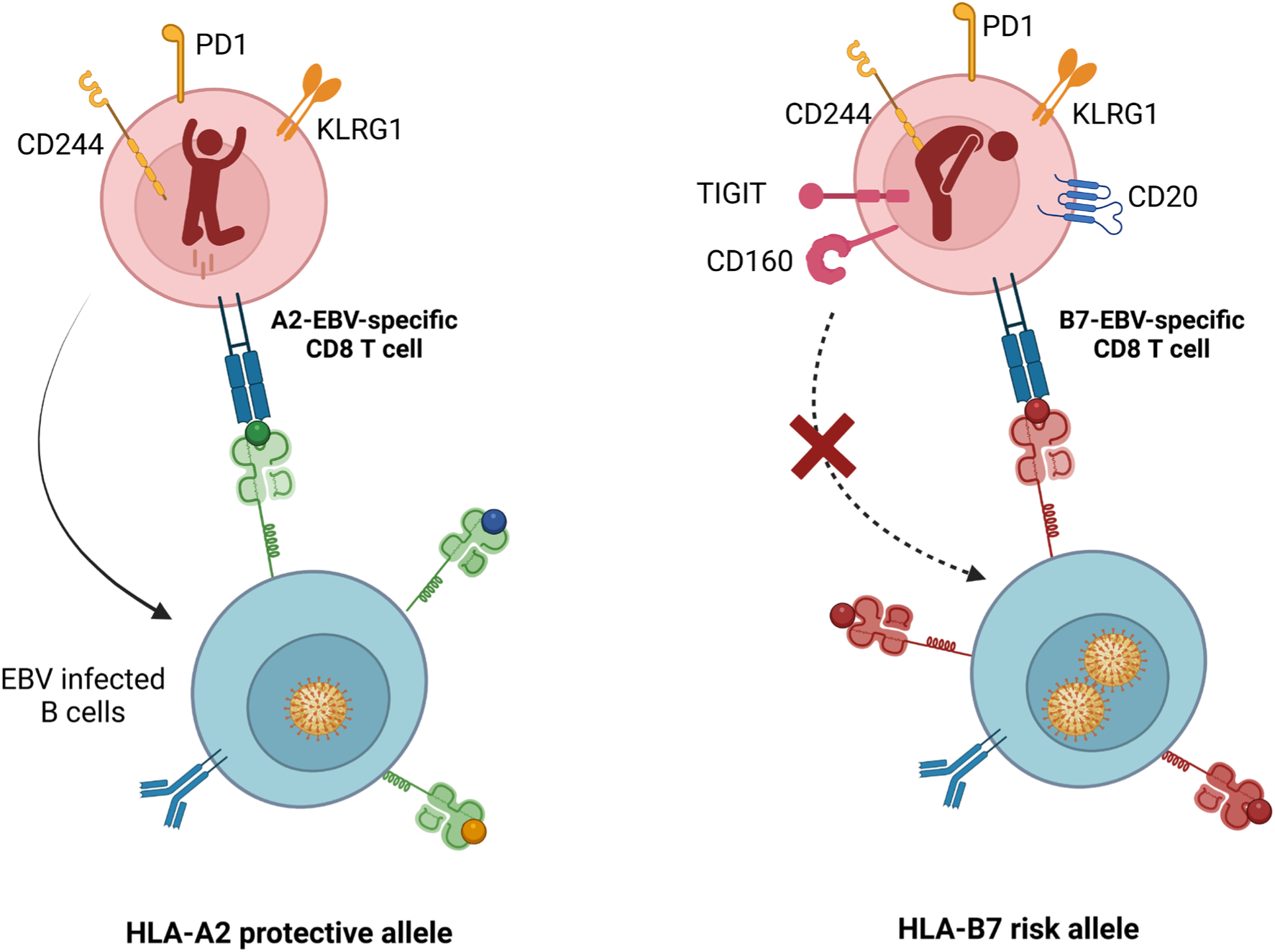
Working model EBV control in MS-associated protective and risk HLA class I alleles. EBV infected B cells with the HLA-A2 protective allele presents a more diverse repertoire of immunodominant EBV peptides compared to the HLA-B7 risk allele. As a consequence, more different CD8+ T cells will recognize EBV when presented in HLA-A2 compared to HLA-B7. For HLA-B7, this could lead to repeated antigen exposure and exhaustion, causing a dysregulation in the control of EBV-infected B cells. Created in https://BioRender.com

## Discussion

The susceptibility to develop MS is multifactorial and includes many genetic and environmental risk factors. In this study, we focused on two prominent risk factors: EBV infection and HLA. We investigated the connection between these major risk factors by investigating EBV-specific CD8^+^ T cells restricted to risk allele HLA-B7 or protective allele HLA-A2. We show that EBV-specific CD8^+^ T cells restricted to risk allele HLA-B7 have a more exhausted phenotype and brain-homing capacity. Since exhausted T cells are less capable of controlling an EBV infection and this might lead to an aberrant immune response, this could explain why HLA-B7 carriers have a higher risk of developing MS compared to HLA-A2 carriers.

Our data show that, for both HD and pwMS, EBV-specific CD8^+^ T cell restricted to HLA-B7 recognized almost solely the latent peptide EBNA3A, while the EBV-specific CD8^+^ T cell restricted to HLA-A2 recognized a broader repertoire of lytic and latent peptides. This is in line with previous findings in HD where it has been shown that frequencies of A2-BMLF1-specific CD8^+^ T cells remain high in the latent phase of an EBV infection, while the frequencies of CD8^+^ T cells specific for other EBV-derived lytic peptides decline (*20*). In the current study, EBV-specific CD8^+^ T cell frequencies were not different between HLA-A2 and HLA-B7 in HD and pwMS. This is variably shown in previous studies, where no differences, higher or lower frequencies were reported (*12, 21, 22*).

In contrast to the similar frequencies, we did find phenotypic differences between CD8^+^ T cells recognizing EBV-derived lytic or latent peptides and differences between EBV-specific CD8^+^ T cells recognizing peptides presented by the risk or protective HLA alleles. In a previous single cell RNA-seq study, a higher expression of CD244 and TIGIT were found in HLA-A2 restricted CD8^+^ T cells specific for EBV-derived lytic versus latent peptides in HD (*11*), which is in line with our extended data on protein level. We report that EBV-specific CD8^+^ T cells restricted to HLA-B7 (EBNA3A) likewise express more co-inhibitory receptors than the ones restricted to HLA-A2 for EBV-derived latent peptides. This difference was even more pronounced in pwMS than HD. The percentage of cells with this exhaustion phenotype was comparable between B7-EBVlatent specific CD8^+^ T cells and the HLA-A2-restricted CD8^+^ T cells specific for EBV-lytic peptides, implying that EBV-specific CD8^+^ T cells restricted to HLA-B7 are exhausted during the latent phase of an EBV infection, while this only occurs during the lytic phase for EBV-specific CD8^+^ T cells restricted to HLA-A2. This implies that the equilibrium between a persistent EBV infection and EBV-specific CD8^+^ T cells is likely disrupted in pwMS during the latent phase. This is relevant as the disease often develops years after primary infection (*6*). Several reports support this interpretation of our data. Jilek et al. reported that, compared to HD, HLA-B7 restricted EBV-specific CD8^+^ T cells in pwMS secrete less IL-2 and IFN-γ after peptide stimulation, which is a functional hallmark of exhausted cells (*12*). We show that the exhaustion phenotype is different between HLA-A2 and HLA-B7 in HD, likely contributing to the increased risk of MS in HLA-B7 carriers. However, a lower cytokine expression in pwMS compared to HD has also been found for the protective allele HLA-A2 (*21*), suggesting that more factors than HLA play a role in the impaired EBV-specific CD8^+^ T cell response in pwMS. Furthermore, Agostini et al. reported that a high EBV viral load in blood of pwMS is highly associated with HLA-B7 (*13*). Reduced clearance of EBV by exhausted CD8^+^ T cells against EBV in HLA-B7^+^ individuals could lead to a higher EBV load. A subsequent increased antigen exposure to EBV-specific B cells could induce a higher antibody titer, which is one of the hallmarks of MS. In line with this hypothesis, we found higher anti-EBNA1 titers in HLA-B7^+^ pwMS.

Next to the exhaustion phenotype, we investigated if HLA also plays a role for the EBV-specific CD8^+^ T cells found in CNS compartments. Several studies have shown an enrichment of EBV-specific CD8^+^ T cells in the CSF and presence of these cells in the brain, which was confirmed in our study (*11, 23–26*). We detected mainly HLA-B7-restricted EBV-specific CD8^+^ T cells in the brain, while Serafini et al. showed similar frequencies of HLA-A2 and HLA-B7-restricted EBV-specific CD8^+^ T cells (*25*). This might be explained by sample variation or differences in used techniques. Serafini et al. used *in situ* staining which is a local visualization, while in our study the mononuclear cells were isolated and analyzed with spectral flow cytometry. We showed that specifically HLA-B7-restricted EBV-specific CD8^+^ T cells express brain residency-associated marker CD20 (*15, 16*) and that this expression remained high on these cells in the different CNS compartments, together with the expression of CXCR6. This phenotype of T cells in the brain is previously described and suggests a higher cytotoxic potential (*15*). A high cytotoxic potential of EBV-specific CD8^+^ T cells in the brain was previously suggested (*25*), highly contrasting with the exhaustion profile found in circulating HLA-B7-restricted EBV-specific CD8^+^ T cells. However, the exhaustion profile is far less prominent in the brain compartments compared to the blood. This might be explained by association of anti-EBNA1 antibodies with susceptibility of developing MS, while the association with MS severity remains controversial (*6, 27*). Genetic SNPs in loci associated with HLA and adaptive immune responses likewise robustly associate with MS risk, while the influence of these factors on MS severity is uncertain (*28*). The reduced EBV control in the circulation for HLA-B7^+^ individuals we currently report, could potentially be more important for susceptibility to MS than for control of EBV in the CNS in relationship with MS pathology.

There are a few limitations in this study. First, the number of subjects was limited due to the strict selection criteria and availability of the samples. All HD and people with MS needed to be selected first on HLA-A2 and HLA-B7. Furthermore, since CD8^+^ EBV-specific T cells are rare and the dominant antigen differs per person, not every sample and specificity reached our cut-off of 20 cells to continue phenotypical analysis. Therefore, we were unable to investigate the phenotype of HLA-B7 restricted CD8^+^ T cells specific for EBV lytic peptides. It would be interesting to investigate if the exhaustion phenotype is even more prominent for these cells. For the CNS compartments, we were able to include one donor. This has to be considered as an anecdotal case and should be further validated. Secondly, we found an exhausted phenotype of HLA-B7-restricted CD8^+^ EBV-specific T cells. We did not perform functional assays, but our phenotype fit with reduced cytokine expression which has been published for HLA-B7-restricted EBV-specific CD8^+^ T cells (*12*). Furthermore, as this is a cross-sectional study, we cannot draw conclusions about the CD8^+^ EBV-specific T cells in the course of the disease. Lastly, the brain residency- associated markers found on the EBV-specific CD8^+^ T cells are phenotypic characteristics. We did not directly show migration, but the presence of the CD8^+^ EBV-specific T cells in the different brain compartments indicate that these are able to migrate towards the brain. Although we did not find an accumulation of total EBV-specific CD8+ T cells in the blood after natalizumab therapy, our data showed that mainly EBV-specific T_EM_ cells were restored in the blood after natalizumab therapy, suggesting that these cells migrate towards the brain. Indeed, the vast majority of the CD8^+^T cell in the brain showed the EM phenotype.

Since EBV is instrumental to the development of MS, preventive strategies could be applied targeting the EBV infection prior to MS onset. An important option would be prophylactic EBV vaccines. Although clinical trials have been performed and are planned, the development of these vaccines prove to be challenging (*29, 30*). Since people living with HIV have a reduced risk on developing MS, treatment with highly active anti-retroviral therapy has been hypothesized to prevent MS-onset in high risk populations (*31*). However, because of the relative low incidence of MS, possible negative health-effects of these drugs and their high costs, this hypothesis needs further investigation. Therapeutic strategies targeting EBV could also be an option as a benefit of reducing EBV viral load in people with MS cannot be excluded. Anti-CD20 therapies deplete circulating memory B cells and are highly effective in pwMS. As B cells are the major target cells of EBV in human body, anti-CD20 therapy also depletes EBV, which might be part of the beneficial result (*32*). Another strategy is to adoptively transfer EBV-specific CD8^+^ T cells. Indeed, the first clinical trial with adoptive transfer of *in vitro*-expanded autologous EBV-specific CD8^+^ T cells has been performed with promising results (*33*). However, this strategy will not overcome T cell exhaustion and could maybe be replaced by TCR gene transfer therapy in the future.

To conclude, our data support the hypothesis that HLA-B7-restricted EBV-specific CD8^+^ T cells are more exhausted and therefore less able to control an EBV infection (*21*). Furthermore, these cells are able to cross the blood-brain barrier in pwMS as these cells were found in the brain. This could be key for the susceptibility to develop MS.

## Materials and methods

### Study design, cohorts and selection criteria

The cohorts used for this study consisted of 58 healthy donors (HD), 419 participants of the PROUD study (UT-MS) and 56 persons with RRMS treated with natalizumab (NTZ-MS). The PROUD study includes persons after their first demyelinating event. In this study, only the participants from the PROUD cohort with a RRMS diagnosis according to the McDonald 2017 criteria (*34*) were included (n=241). HLA-A2^+^ and/or HLA-B7^+^ individuals were selected from these 3 cohorts (17 HD, 15 untreated pwMS and 15 NTZ-treated pwMS) for spectral flow cytometry. For all the selected participants from the NTZ-MS cohort, the time of blood sampling was between 6 and 18 months after the first round of treatment. The gender, age and HLA alleles of selected participants for the 3 cohorts are described in table S1. The Netherlands Brain Bank donor was a HLA-A2^+^B7^+^ male of 77 years old with MS. The PROUD study, ErasMS biobank and collection of the HD material were approved by the ethics committee of Erasmus MC, University Medical Center Rotterdam (MEC-2021-0946, MEC-2022-0389, MEC-2021-0251, respectively). Informed consent forms were signed by all participants. The Netherlands Brain Bank donor program was approved by the ethics committee of the Vrije Universiteit Medical Center, Amsterdam (2009/148).

### Isolation of mononuclear cells from peripheral blood

Peripheral blood was collected in CPT tubes, the PBMCs were isolated according to manufacturer’s protocol and stored in liquid nitrogen. Serum was collected and stored at −80 freezer. Peripheral blood from the MS brain donor was obtained post-mortem via cardiac puncture and PBMCs were isolated using Ficoll-Plaque Plus (GE Heathcare) as previously described (*8*).

### Isolation of mononuclear cells from post-mortem CNS compartments

Cerebrospinal fluid (CSF), leptomeninges (LM), normal appearing white matter (NAWM) and lesion from a person with MS were derived from the Netherlands Brain Bank. Mononuclear cells were isolated from the different compartments as previously described (*8, 35, 36*), with the addition of erythrocyte lysis and a modification for the enzymatic incubation step. Per 2-3 gram LM, NAWM or lesion, 7.5mg Collagenase type IV (Worthington) together with 500U DNAse I rec (Roche) in DPBS+Ca/Mg+0.1% BSA were added and incubated at 37°C for 60 min before mechanical dissociation.

### HLA genotyping

Genomic DNA from the PROUD cohort was isolated from blood samples collected at study baseline. Samples were genotyped using Illumina Infinium High-Throughput Screening (HTS) iSelect custom-730K SNP beadchip array. Next, 4-digit HLA typing was performed by HLA imputation using the HLA-TAPAS algorithm (*37*). For the HD cohort, NTZ-MS cohort and brain donor, genotyping for HLA-A2 and HLA-B7 was performed by flow cytometry. PBMCs were stained with HLA-A2-PE and HLA-B7-APC for 15 min at RT in the dark, washed with FACS buffer (PBS+0.5%BSA+0.01% Sodium Azide) and measured on a BD LSRFortessa Flow Cytometer (BD Biosciences). The antibodies are listed in table S2.

### Peptides

A selection of the most immunodominant peptides based on the immune epitope database (iedb.org) (*38*) were ordered at JPT peptide technologies (>70% purity, N-terminus amine, C-terminus acid, counter-ion trifluoroacetate), dissolved in MilliQ to a concentration of 1mg/ml and stored at −20° C till further use. All peptides are listed in table S3.

### EBV- and CMV-specific tetramers

EBV- and CMV- specific cells were identified using in house prepared HLA-A*02:01 and HLA-B*07:02 tetramers (*39*), making use of monomers with photocleavable peptides (*40, 41*). EBV- and CMV-derived peptides were exchanged as previously described (*42*). In short, the monomers and peptides were diluted in exchange buffer (20mM Tris + 100mM NaCl, pH=8.0), mixed and exposed for 1 hour to UV light. A separate tube with monomers was used for every peptide. Next, fluorochrome-coupled streptavidin (Table S2) diluted in exchange buffer was added in 10 steps with 15min incubations on RT in between. Every peptide-monomer combination was separately labeled with 2 different fluorochromes. Tetramers were stored at 4° C.

### Flow cytometry

Frozen PBMCS were thawed, counted and 5×10^6 PBMCs were used per staining. First, the cells were stained with Zombie NIR Fixable Viability Kit (Biolegend) for 10min at RT. Cells were washed once with FACS buffer and resuspended in Brilliant Stain Buffer (BSB; BD Biosciences). Fc Receptor blocking solution, CCR7 and CCR2 were added and incubated 5 min at RT, followed by CCR6 2 min at RT, CXCR6 and CCR4 2 min at RT, CCR5 2 min at RT and an incubation of 20 min at RT after adding CXCR3. The cells were washed twice with FACS buffer and stained with the HLA-A2 or HLA-B7 tetramer mix. For the HLA-A2^+^B7^+^ samples, 2 separate protocols were performed with each 5×10^6 cells, one using tetramers for HLA-A2 and one for HLA-B7. After 15 min incubation at RT, cells were washed twice with FACS buffer. Subsequently, the following antibodies were added 1 by 1 in BSB and incubated 20 min at RT: CD28, GPR56, CD69, CD20, CD127, SLAMF7, CX3CR1 and CD29. Cells were washed twice with FACS buffer and stained with an antibody mix diluted in FACS buffer: CD4, CD161, CD19, CD8, CD160, CD3, CD56, CD16, CD244, KLRG1, CD26, PD-1, CD45RA, CD38, TIGIT and CD103. After 20 min incubation at RT, cells were washed twice in FACS buffer and resuspended in PBS+0.1% BSA. The samples were measured on a Cytek Aurora 5 laser spectral cytometer. All antibodies are listed in table S2.

### Serology

Serum samples from participants were collected at baseline for the PROUD study and 6 months to 7 years after treatment for the NTZ-MS cohort. Epstein-Barr virus (EBV) nuclear antigen EBNA-1 (NA), early antigen (EA) and viral capsid antigens (VCA) and Cytomegalovirus (CMV) IgG antibodies were measured using the LIAISON® IgG chemiluminescent immunoassays (CLIA) on the LIAISON® XL analyzer (DiaSorin, Italy). The serological test and the analysis were conducted according to the manufacturer’s instructions. The serum samples were 20x diluted to prevent results from exceeding the detection limit of the tests. To ensure reliability, quality control samples supplied by the manufacturer were included in each testing batch.

### Data analysis

Flow cytometry data were analyzed using OMIQ software form Dotmatics. EBV- and CMV specific T cells were gated manually and later pooled as EBV-latent, EBV-lytic or CMV-specific cells. Reference PBMCs from the same donor were taken along in every experiment to adjust for technical variation in marker expression. The expression level of the markers in the samples were normalized with CytoNorm using the reference PBMCs (*43*). Samples with fewer than 20 antigen-specific cells were excluded from the analysis. For the brain compartments, CX3CR1 was used to exclude microglia.

### Statistical analysis

Statistical analysis was done in Graphpad Prism 9.2.0. The most suitable statistical tests were chosen based on the distribution and variance in the data and are described in the figure legend. A P value lower than 0.05 was considered significant.

## Acknowledgements

We thank Jard de Vries from the department of Internal Medicine of the Erasmus MC for the HLA imputation, Dr. Corine H. Geurts van Kessel for supervising serological testing at the department of Viroscience of the Erasmus MC for the serology, and Peter van Geel for assistance at the Flow cytometry shared facility of the Erasmus MC.

## Funding

Stichting MS research, fellowship grant (21-1142), MvL

Stichting MS research, onderzoeksprogramma (23-490g), JS

Nationaal MS Fonds (OZ2021-016), MvL

Nationaal MS Fonds, educational grant (P2021-001), CC

MoveS stichting klimmen tegen MS, Inspiratiebeurs, JS

European Union’s Horizon Europe Research and Innovation Actions, BEHIND-MS (101137235), MvL

## Author contributions

Conceptualization: SR, RN, JS, MvL

Investigation: SR, JR, AM, AW, HW, MJM, VM, JvL

Visualization: SR, AM, JR

Statistical analysis: SR, YvH

Resources: CC, YM, BW

Funding acquisition: CC, JS, MvL

Supervision: JS, MvL

Writing – original draft: SR

Writing – review & editing: SR, JR, AM, AW, MJM, CC, YvH, YM, JS, MvL

## Data and materials availability

Data will be made available by the corresponding author upon reasonable request by qualified researchers. For sharing individualized data or material of participants who provided consent for sharing, a data transfer agreement (DTA) or material transfer agreement (MTA) needs to be developed as appropriate.

