## Supplemental file for "Multiple sclerosis-associated HLA demarcates EBV-specific CD8^+^ T cells with an exhausted and brain residency phenotype"

**Table S1. Characteristics of selected samples for flow cytometry**

| | **HD (n=17)** | **UT-MS (n=15)** | **NTZ-MS (n=15)** |
|---|---|---|---|
| **Sex** | | | |
| Female, n (%) | 9 (53) | 9 (60) | 12 (80) |
| Male, n (%) | 8 (47) | 6 (40) | 3 (20) |
| **Age** | | | |
| Median | 43 | 30 | 38 |
| Range | 25-69 | 25-50 | 23-62 |
| **HLA** | | | |
| $A2^{+}B7^{-}$ | 6 | 5 | 4 |
| $A2^{-}B7^{+}$ | 7 | 5 | 4 |
| $A2^{+}B7^{+}$ | 4 | 5 | 7 |

**Table S2. Antibodies**

| Target | Fluorochrome | Clone | Company | Catalog number |
|---|---|---|---|---|
| HLA-A2 | PE | BB7.2 | BD Biosciences | 558570 |
| HLA-B7 | APC | BB7.1 | Biolegend | 372405 |
| CD28 | BUV496 | CD28.2 | BD Biosciences | 741168 |
| GPR56 | BUV563 | CG4 | BD Biosciences | 752709 |
| CD69 | BUV737 | RN50 | BD Biosciences | 612818 |
| CD20 | BUV805 | 2H7 | BD Biosciences | 612905 |
| CD196/CCR6 | BV421 | G034E3 | Biolegend | 353408 |
| CD161 | Pacific Blue | HP-3G10 | Biolegend | 339926 |
| CD192/CCR2 | BV480 | 1D9 | BD Biosciences | 747852 |
| CD19 | eFluor506 | HIB19 | ThermoFisher | 69-0199-42 |
| CD8 | cFluor V547 | SK1 | Cytek | SKU R7-20063 |
| CD127 | BV570 | A019D5 | Biolegend | 351308 |
| CD194/CCR4 | BV605 | L291H4 | Biolegend | 359417 |
| CX3CR1 | BV650 | 2A9-1 | Biolegend | 341626 |
| CD195/CCR5 | BV711 | 2D7 | BD biosciences | 563395 |
| CD29 | BV750 | MAR4 | BD Biosciences | 747231 |
| CD197/CCR7 | BV785 | G043H7 | Biolegend | 353230 |
| CD160 | AF488 | BY55 | BD biosciences | 562351 |
| CD3 | AF532 | UCHT1 | ThermoFisher | 58-0038-41 |
| CD16 | NovaFluor 610-70S | 3G8 | ThermoFisher | H006T02B06 |
| CD319/SLAMF7 | BB700 | 235614 | BD Biosciences | 749692 |
| CD56 | PerCP-eFluor710 | TULY56 | ThermoFisher | 46-0566-42 |
| CD103 | RB780 | BER-ACT8 | BD Biosciences | 569331 |
| CD186/CXCR6 | RY586 | 13B 1E5 | BD biosciences | 753575 |
| CD183/CXCR3 | PE-Dazzle594 | G025H7 | Biolegend | 353736 |
| PD-1 | PE-Cy5 | EH12.2H7 | Biolegend | 329971 |
| CD244/2B4 | PE-Cy5.5 | C1.7 | ThermoFisher | 35-5838-42 |
| KLRG1 | PE-Fire810 | SA231A2 | Biolegend | 367733 |
| CD4 | AF647 | SK3 | BD Biosciences | 344635 |
| CD45RA | SparkNIR685 | HI100 | Biolegend | 304167 |
| CD26 | AF700 | 222113 | R&D Systems | FAB1180N-100ug |
| TIGIT | APC-Cy7 | A15253G | Biolegend | 372733 |
| CD38 | APC-Fire810 | HIT2 | Biolegend | 303550 |
| Fixable Viability Kit | Zombie NIR | n/a | Biolegend | 423106 |
| Streptavidin | BUV395 | n/a | BD Biosciences | 564176 |
| Streptavidin | BUV661 | n/a | BD Biosciences | 612979 |
| Streptavidin | PE | n/a | ThermoFisher | S866 |
| Streptavidin | PE-Cy7 | n/a | Biolegend | 405206 |
| Streptavidin | APC | n/a | ThermoFisher | SA1005 |

| | | | | |
|---|---|---|---|---|
| Human TruStain FcX | n/a | n/a | Biolegend | 422302 |

**Table S3. Peptides**

| **Peptide** | **Sequence** | **HLA restriction** | **Virus** | **Lytic or latent protein** |
|---|---|---|---|---|
| BMLF1(280-288) | GLCTLVAML | HLA-A*02:01 | EBV | Lytic |
| BRLF1(109-117) | YVLDHLIVV | HLA-A*02:01 | EBV | Lytic |
| BMRF1(208-216) | TLDYKPLSV | HLA-A*02:01 | EBV | Lytic |
| BALF4(276-284) | FLDKGTYTL | HLA-A*02:01 | EBV | Lytic |
| LMP2(356-364) | FLYALALLL | HLA-A*02:01 | EBV | Latent |
| LMP2(426-434) | CLGGLLTMV | HLA-A*02:01 | EBV | Latent |
| LMP1(125-133) | YLLEMLWRL | HLA-A*02:01 | EBV | Latent |
| EBNA3C(284-293) | LLDFVRFMGV | HLA-A*02:01 | EBV | Latent |
| pp65(495-503) | NLVPMVATV | HLA-A*02:01 | CMV | Lytic |
| IE(316-324) | VLEETSVML | HLA-A*02:01 | CMV | Lytic |
| BMRF1(116-128) | RPQGGSRPEFVKL | HLA-B*07:02 | EBV | Lytic |
| BZLF1(44-52) | LPCVLWPVL | HLA-B*07:02 | EBV | Lytic |
| EBNA3A(379-387) | RPPIFIRRL | HLA-B*07:02 | EBV | Latent |
| EBNA3C(881-889) | QPRAPIRPI | HLA-B*07:02 | EBV | Latent |
| EBNA1(528-536) | IPQCRLTPL | HLA-B*07:02 | EBV | Latent |
| pp65(265-275) | RPHERNGFTVL | HLA-B*07:02 | CMV | Lytic |
| pp65(417-425) | TPRVTGGGAM | HLA-B*07:02 | CMV | Lytic |
| IE(309-317) | CRVLCCYVL | HLA-B*07:02 | CMV | Lytic |

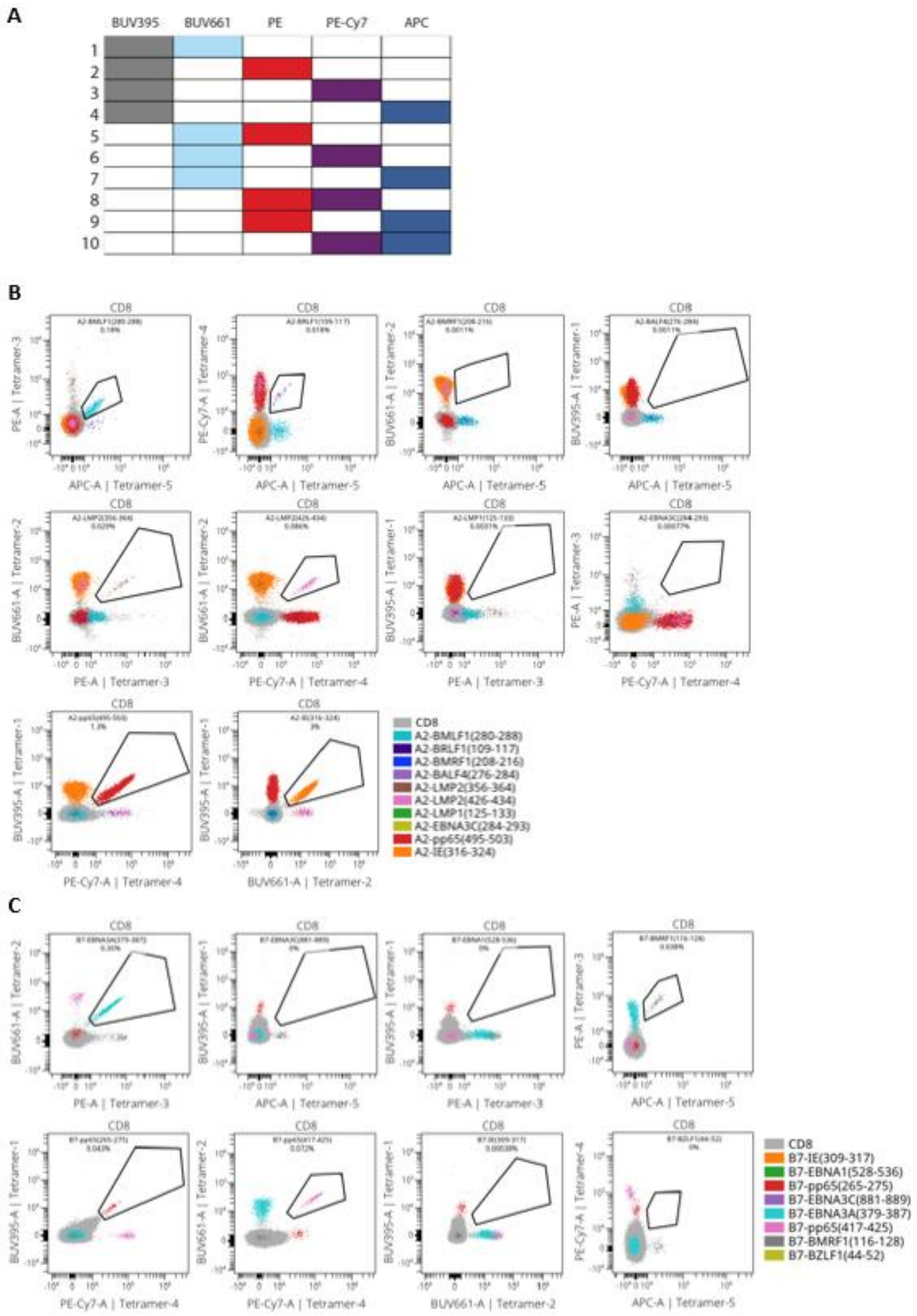


**Figure S1. EBV- and CMV-specific CD8$^{+}$ T cell gating. (A)** Combinatorial staining method for HLA class I tetramers. The columns depict the 5 different fluorochromes used to conjugate tetramers. Each row depict an antigen-specific tetramer. **(B)** Representative example tetramer staining on a HD with HLA-A2 EBV and CMV tetramers. **(C)** Representative example tetramer staining on a HD with HLA-B7 EBV and CMV tetramers.

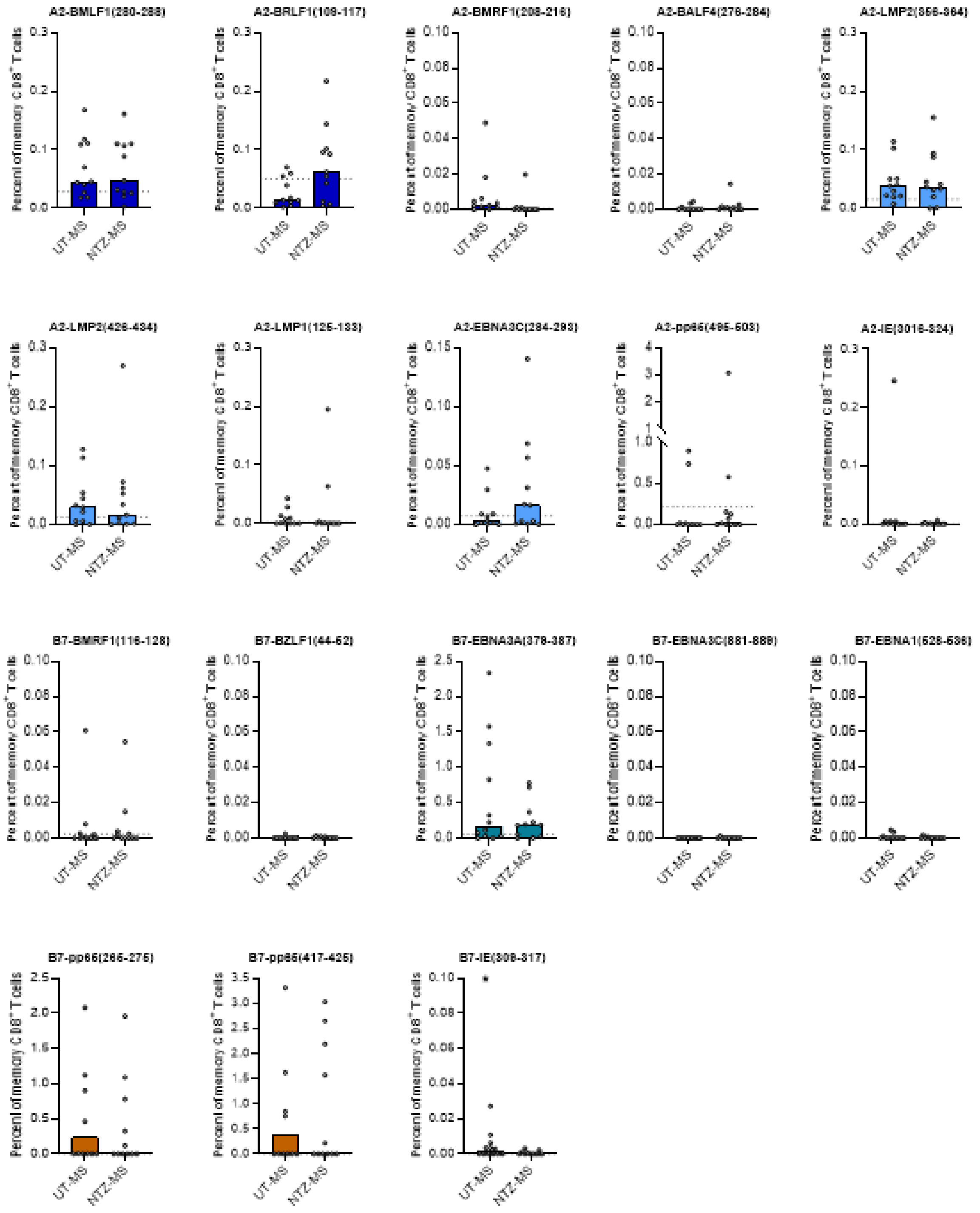


**Figure S2. Frequencies of EBV- and CMV-specific CD8+ T cells in MS for every epitope.** Every dot represents one individual. Bars represent medians. Dotted lines represents median of HD. Mann-Whitney tests were used for statistical testing *p<0.05 ** p<0.005 ***p<0.0005 **** p<0.0001.

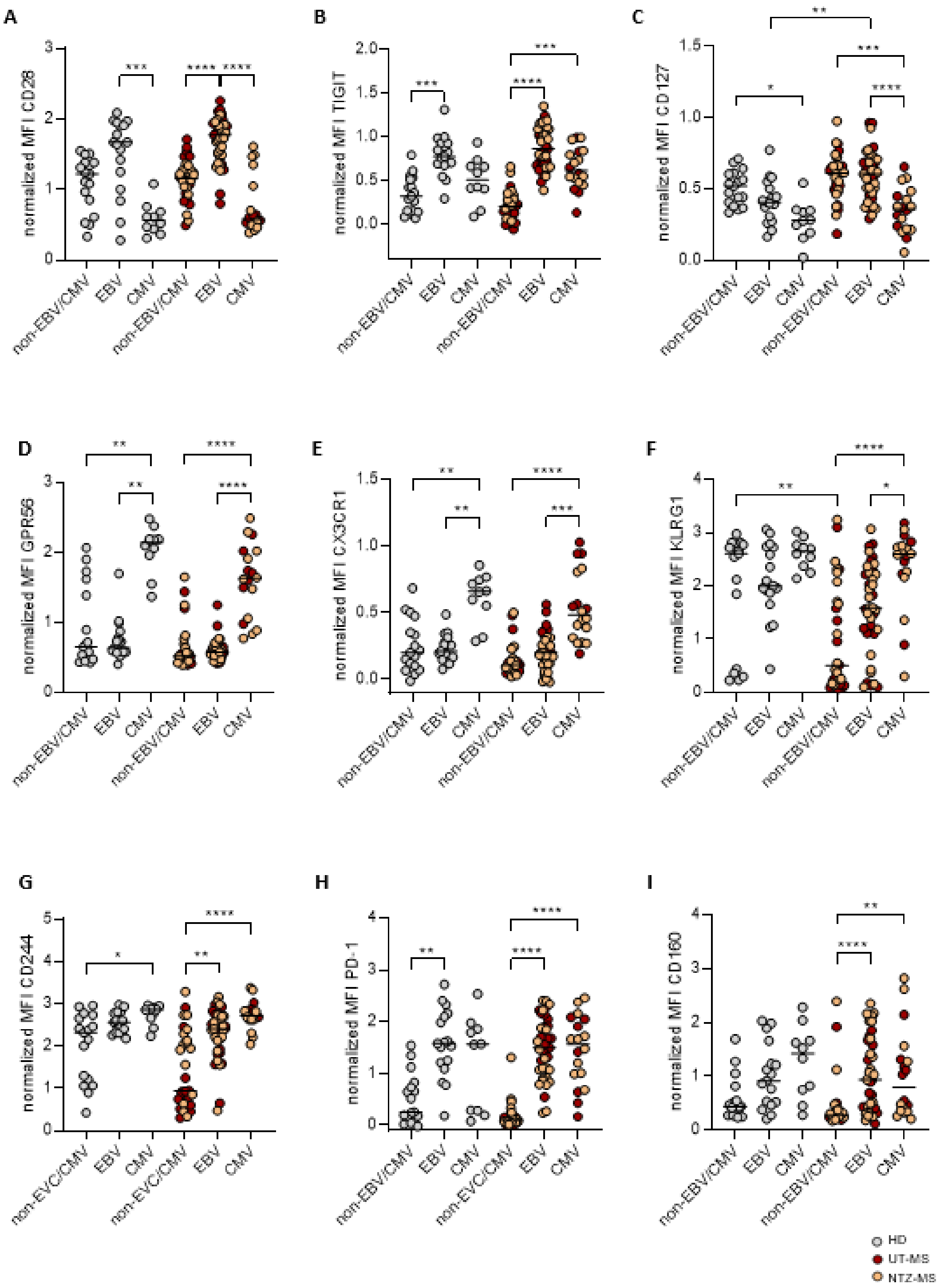


**Figure S3. Differential marker expression between EBV- and CMV-specific CD8+ T cells.** Median Fluorescence Intensity (MFI) of **(A)** CD28 **(B)** TIGIT **(C)** CD127 **(D)** GPR56 **(E)** CX3CR1 **(F)** KLRG1 **(G)** CD244 **(H)** PD-1 and **(I)** CD160 in EBV-specific, CMV-specific and total CD8+ memory T cells. Kruskal-Wallis test with Dunn posthoc was used for statistical testing. Every dot represents one individual. Grey dots are HD, red dots are UT-MS and orange dots are NTZ-MS. Black lines represent medians. *p<0.05 ** p<0.005 ***p<0.0005 **** p<0.0001.

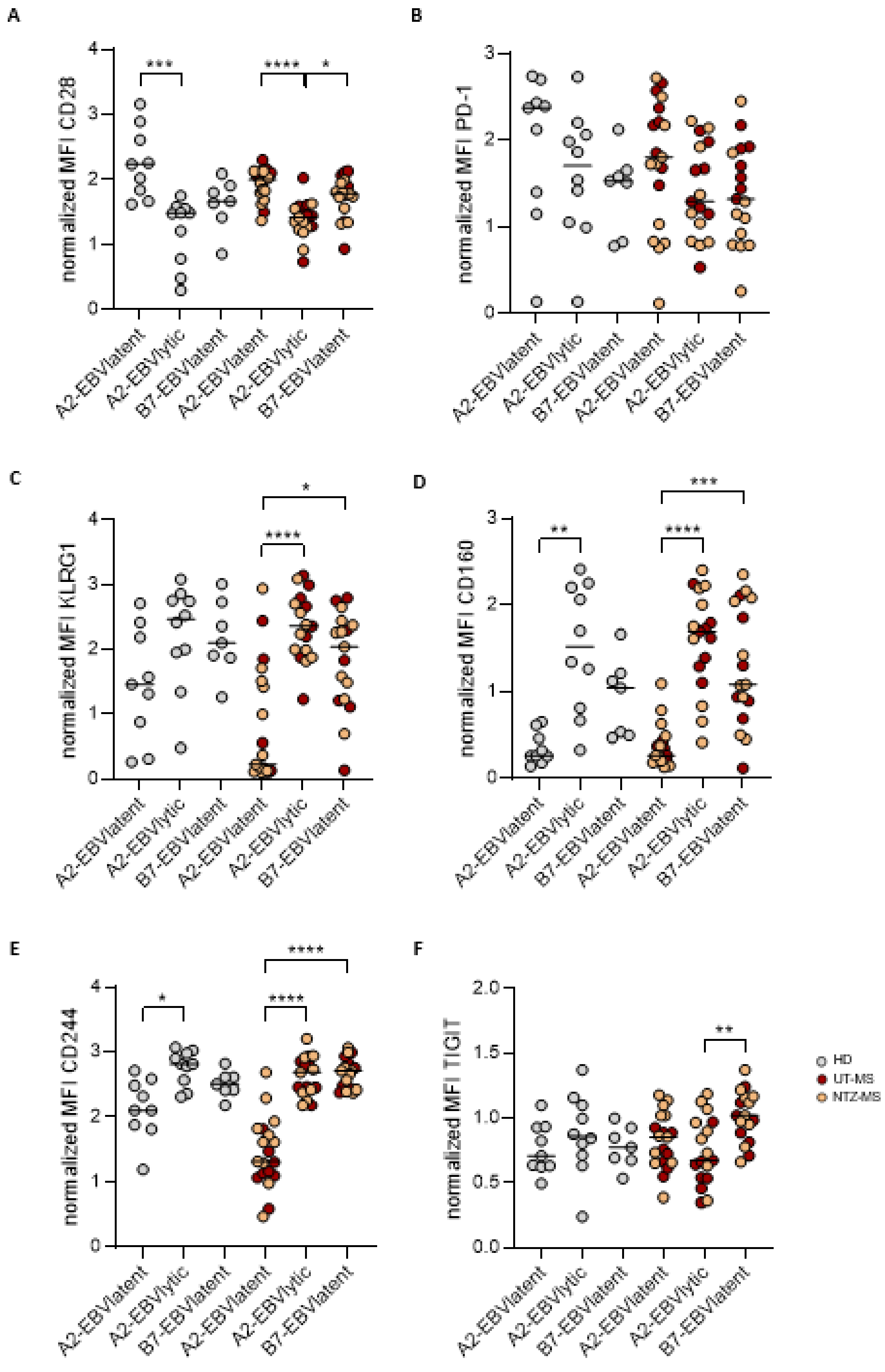


**Figure S4. Co-stimulatory and co-inhibitory marker expression on EBV-specific CD8+ T cells recognizing lytic or latent epitopes.** Median fluorescence intensity (MFI) of **(A)** CD28 **(B)** PD-1 **(C)** KLRG1 **(D)** CD160 **(E)** CD244 and **(F)** TIGIT. Kruskal-Wallis test with Dunn posthoc was used for statistical testing. Every dot represents one individual. Grey dots are HD, red dots UT-MS and orange dots are NTZ-MS. Black lines represent medians. *$p<0.05$ ** $p<0.005$ ***$p<0.0005$ **** $p<0.0001$.

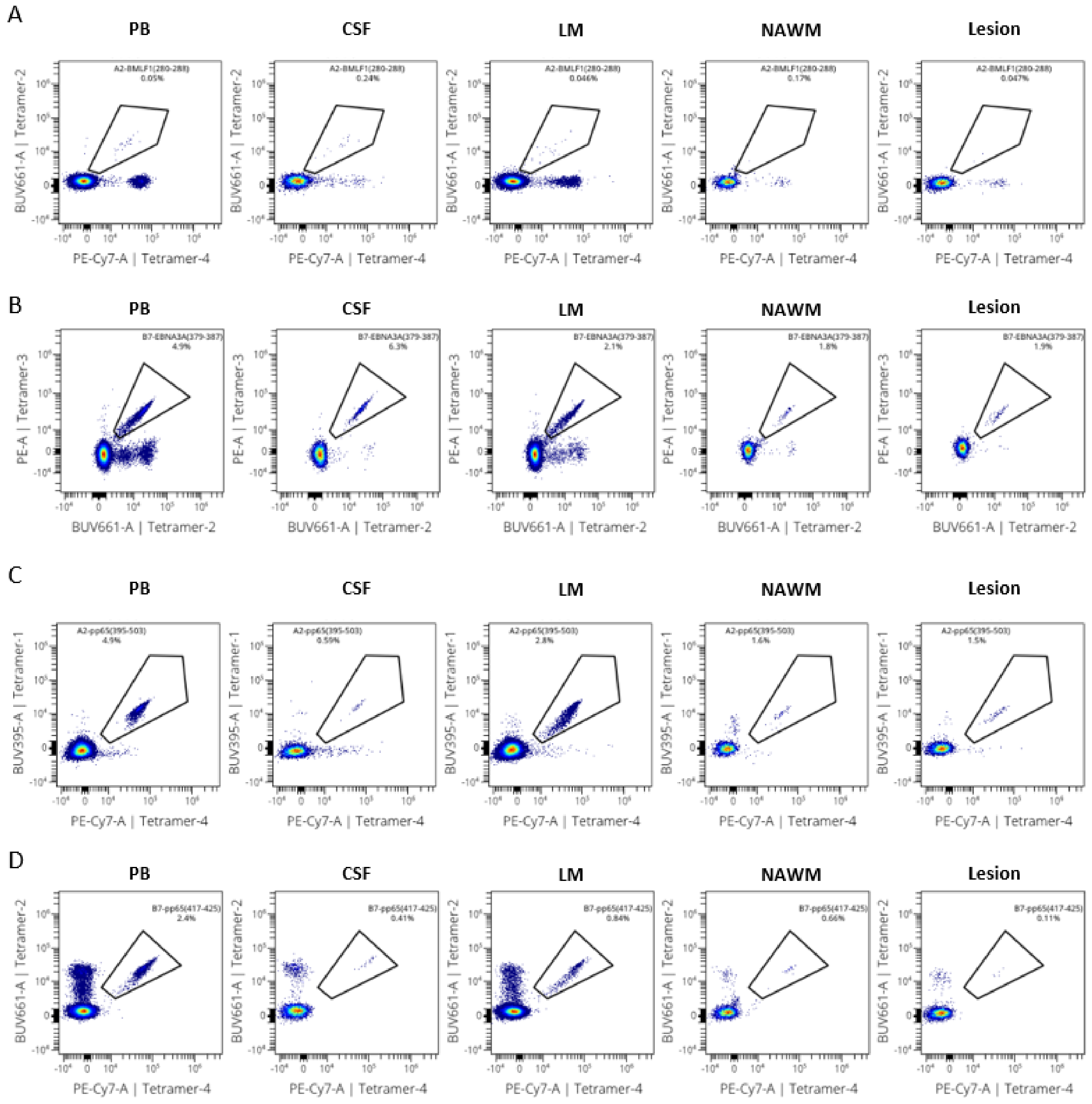


**Figure S5. Gating of EBV- and CMV-specific CD8⁺ T cells in different post-mortem CNS compartments of an MS donor. (A)** EBV A2-BMLF1(280-288) **(B)** EBV B7-EBNA3A(379-387) **(C)** CMV A2-pp65(495-503) **(D)** CMV B7-pp65(417-425).

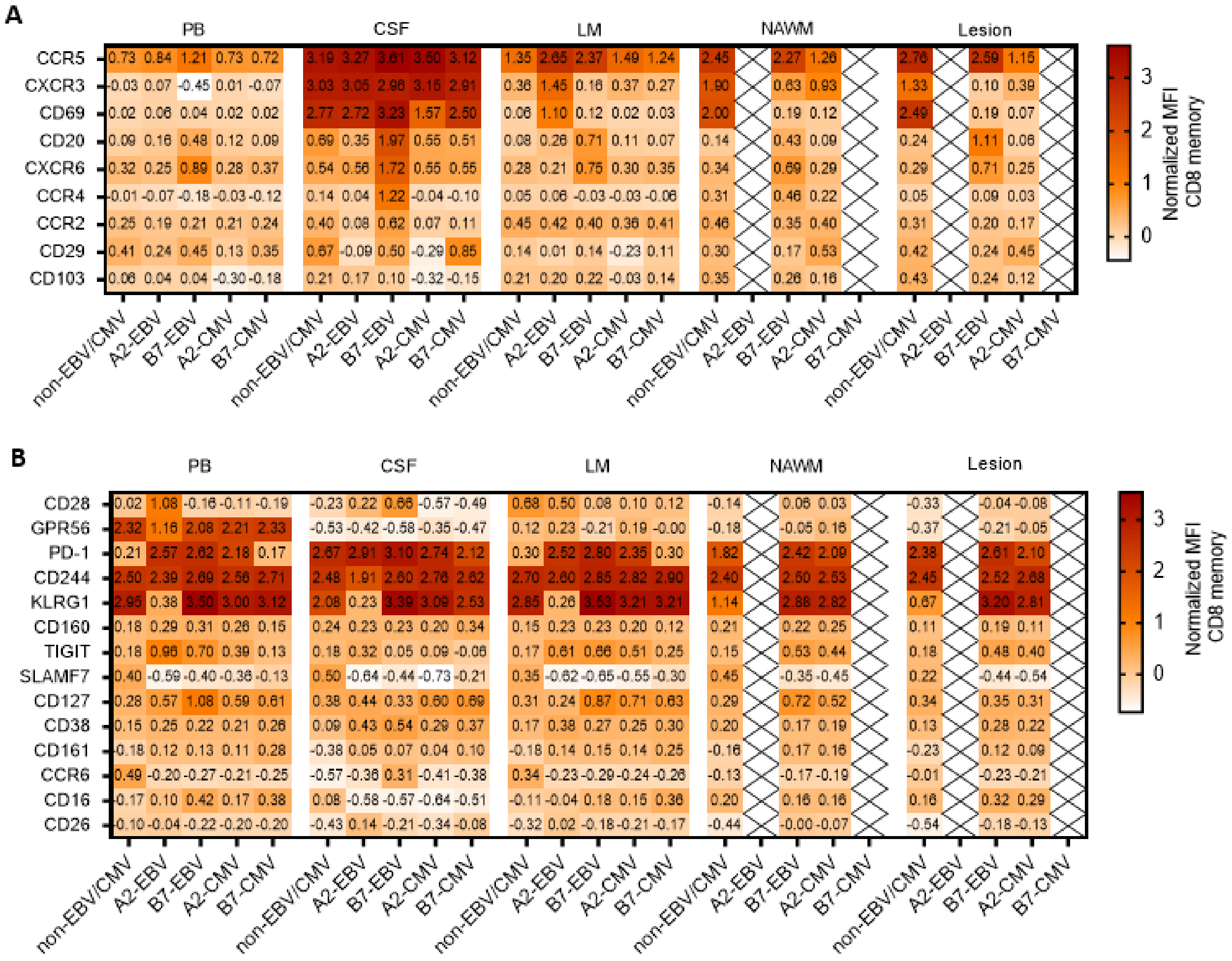


**Figure S6. Brain-homing/residency- and activation-associated markers in CNS compartments.** Heatmap of median protein expression in normalized MFI for total CD8$^+$ memory T cells, EBV- and CMV-specific CD8$^+$ memory T cells for **(A)** brain-homing/residency-associated markers or **(B)** activation-associated markers. Data from 1 brain donor with MS.

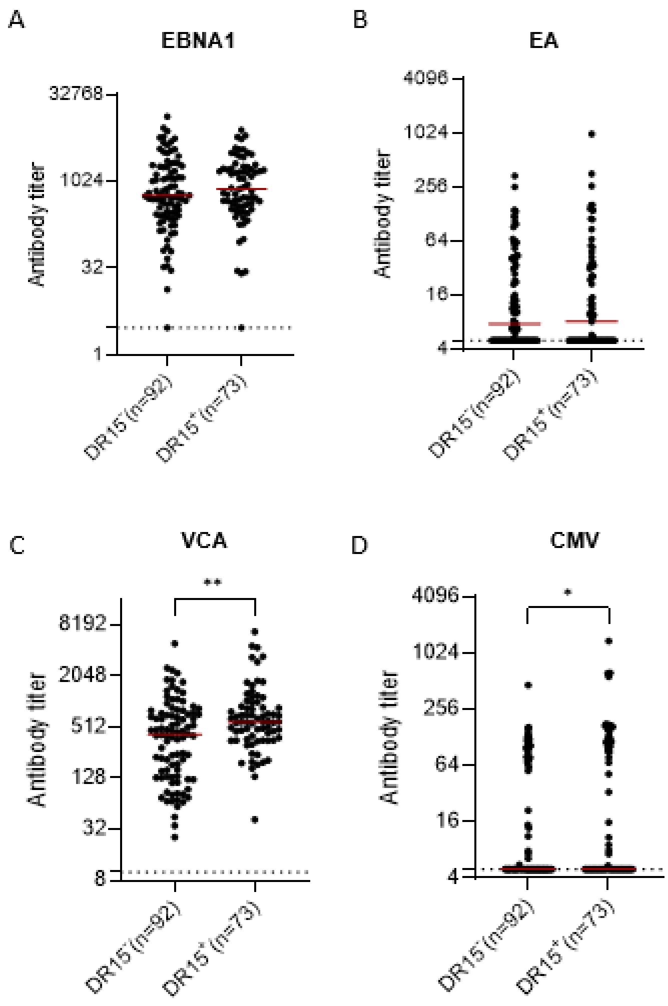


**Figure S7. Antibody titers in untreated pwMS stratified on HLA-DRB1*15:01.** Antibody titers of HLA-DRB1*15:01$^{-}$ or HLA-DRB1*15:01$^{+}$ untreated persons with RRMS from the PROUD cohort. **(A)** EBNA1 **(B)** Early antigen (EA) **(C)** Viral capsid antigen (VCA) **(D)** CMV. Mann Whitney test was used for statistical testing. Red lines represent medians with 95% CI. *p<0.05 ** p<0.005 ***p<0.0005 **** p<0.0001.
